# Busy and confused? High risk of missed alerts in the cockpit: an electrophysiological study

**DOI:** 10.1101/2022.01.07.475323

**Authors:** Mickael Causse, Fabrice B. R. Parmentier, Damien Mouratille, Dorothée Thibaut, Marie Kisselenko, Eve Fabre

## Abstract

Of evolutionary importance, the ability to react to unexpected auditory stimuli remains critical today, especially in settings such as aircraft cockpits or air traffic control towers, characterized by high mental and auditory loads. Evidences show that both factors can negatively impact auditory attention and prevent appropriate reactions in hazardous situations. In the present study, sixty participants performed a simulated aviation task, varying in terms of mental load (no, low, high mental load), that was embedded with a concurrent tone detection paradigm, in which auditory load was manipulated by the number of different tones (1, 2 or 3). We measured both detection performance (miss, false alarm) and brain activity (event-related potentials) related to the target tone. Our results showed that both mental and auditory loads affected tone detection performance. Importantly, their combined effects had a massive impact on the percentage of missed target tones. While, in the no mental load condition, miss rate was very low with 1 (0.53%) and 2 tones (1.11%), it increased drastically with 3 tones (24.44%), and this effect was accentuated as mental load increased, yielding to the higher miss rate in the 3-tone paradigm under high mental load conditions (68.64%). Increased mental load, auditory load, and miss rate, were all associated with disrupted brain response to the target tone as showed by reductions of the P3b amplitude. In sum, our results highlight the importance of balancing mental and auditory loads to maintain or improve efficient reactions to alarms in complex environment.

## INTRODUCTION

### 1.1. Reacting to auditory alarms in challenging environments

The responsiveness to unpredictable auditory stimuli, in the form of an orienting response (e.g., Sokolov, 1963), has long been identified as a fundamental aspect of our cognitive functioning and is thought to have played an essential role in the survival of human beings across evolution (e.g., to alert us of a potential danger and help us detect and flee an approaching predator). Nowadays, this ability to detect and involuntarily orient our attention toward an unexpected sound while voluntarily attending an ongoing task remains a useful feature of our cognitive system in many everyday life situations. This mechanism enables the detection of potential dangers and modifies our actions, allowing for example the sudden and unexpected sound of an approaching vehicle to break into our focus of attention as we are crossing a road. This is of special relevance in many complex working environments where operators must react promptly and accurately to auditory alarms, such as in aircraft cockpits, control towers, operating rooms, or nuclear power plants to name a few (e.g., Guillaume, 2011). Auditory alarms are effective stimuli for conveying critical information in visually saturated environments. Indeed, their near omnidirectional character (i.e., “gaze-free”) yields a higher probability of being detected compared to visual signals (Van der Heiden et al., 2018), and they do not require an effortful and systematic scan of a control panel (Guillaume, 2011). Accordingly, anesthetists have been found to react faster to auditory alarms than to visual alarms, as the latter are only detected upon voluntary inspection of the information presented on a monitor (Morris & Montano, 1996). For this reason, safety-critical aviation events such as cabin decompression, stall, or overspeed are generally associated with auditory alarms in the cockpit.

To improve safety, operators are required to maintain an up-to-date situational awareness (Stanton et al., 2001). This requires the ability to perform certain tasks while monitoring the environment for unexpected stimuli (Causse, Peysakhovich, et al., 2016) that are not necessarily related to the task at hand, but that may potentially convey important information for safety, for instance, a depressurization alarm sounding while the crew is busy performing a landing maneuver. Unfortunately, many accident reports and experimental studies in aeronautics and other complex domains have shown that auditory alarms can be neglected by operators (Bliss, 2003; Dehais et al., 2014; Giraudet et al., 2012; Van der Heiden et al., 2018). Sometimes, operators detect these alarms but fail to react because they misunderstand them (Dehais et al., 2014), consider them unreliable (Bliss & Acton, 2003; Bliss & Dunn, 2000), or underestimate their urgency (Salas & Schlesinger, 2019). However, on occasions, their absence of reaction results from the failure to detect these alarms in the first place. Growing evidence suggests that the capacity to involuntary detect and process auditory information may be modulated by the level of engagement in the task at hand (Van der Heiden et al., 2018). Selective attention helps focusing on the task at hand and reduces potential distraction, but such filtering can result in missing safety-relevant signals. This phenomenon known as *inattentional deafness* (Giraudet, St-Louis, et al., 2015; Koreimann et al., 2014; Macdonald & Lavie, 2011; Raveh & Lavie, 2015) can dramatically affect safety.

### 1.2. Mental load

Aviation accident reports show that when pilots missed an alarm, they were frequently focusing on another mentally-demanding task (see for instance the accident of Eastern Air Lines Flight 401; NTSB, 1973). Indeed, mentally demanding tasks – characterized by high levels of mental load – can affect both the detection of auditory stimuli and the allocation of attentional resources to the processing of their content (e.g., Causse, Imbert, et al., 2016; SanMiguel et al., 2008). This is in line with the classical proposition of an inverse relationship between mental load and attentional reserve which, when depleted, limits the processing of additional information (Kahneman, 1973). Working memory (WM) limitations have certainly a central role in this attentional bottleneck (Tombu et al., 2011). WM is viewed as a system in charge of maintaining and processing information simultaneously, as well as controlling attention in order to select relevant information and resist interference. Some evidence suggests that while unexpected sounds are normally involuntarily detected (see Parmentier, 2014, for a review), this detection appears to be reduced when the primary task’s demands on WM increases (Berti & Schröger, 2003). In this respect, a high mental load may alter the automatic orienting of attention towards unexpected stimuli.

In this sense, a number of behavioral and neuroscientific studies show that operators are more likely to fail to detect and respond to stimuli associated with a secondary task if the cognitive demands of the primary task increases (for an overview, see Ghani et al., 2020). Various studies have also revealed that task load impacts the balance between primary task engagement and the capacity to remain responsive to auditory stimuli (e.g., Berti & Schröger, 2003; Zhu et al., 2022). In numerous studies, the primary task’s difficulty is manipulated and its impact on the passive or active processing of auditory stimuli is examined via the analysis of event-related potentials (ERPs) (e.g., Ghani et al., 2020; Gibson et al., 2019; Giraudet, Imbert, et al., 2015; Kramer et al., 1995; M. M. Swerdloff & L. J. Hargrove, 2020; Raabe et al., 2005; Ullsperger et al., 2001). The tasks used to evaluate the passive processing of auditory stimuli generally consist in presenting task-irrelevant auditory stimuli (also called distractors), to which no active response is expected from participants. Overall, these studies have shown that an increase in task load negatively affects the processing of the auditory stimuli (e.g., Berti & Schröger, 2003; Fabre et al., 2017; Lv et al., 2010; SanMiguel et al., 2008). For example, using a categorization task, San Miguel et al. (2008) reported a reduction of the distraction caused by novel sounds in a 1-back condition relative to a no load condition. Studies assessing the active processing of auditory stimuli require the participant to detect a target sound and to produce an action in response (e.g., to press a response button), in a way comparable to how operators are expected to behave after the occurrence of an alarm (e.g., Callan et al., 2018; Causse, Imbert, et al., 2016; Dehais, Duprès, et al., 2019; Giraudet, St-Louis, et al., 2015). Participants perform a primary task (e.g., piloting task; Causse, Peysakhovich, et al., 2016; Giraudet, St-Louis, et al., 2015; Kramer et al., 1987) while concurrently doing an alarm detection task (referred to as a secondary task). This detection task can take the form of an oddball paradigm (Squires et al., 1975) in which participants must respond to infrequent target tones embedded in a stimulus train that may also include a varying proportion of other, non-target, sounds.

From a neurological stand point, several authors have suggested that brain responses to the auditory stimuli reliably measure mental load elicited by the primary task (Kramer et al., 1995; Miller et al., 2011). In general, the results show that the detection of the target sounds decreases as workload increases (e.g., Giraudet, St-Louis, et al., 2015), and the workload impact brain electrophysiological responses to target sounds, at different stages of the information processing, as indexed by the N100 and the P300 components (Dehais, Roy, et al., 2019; Giraudet, St-Louis, et al., 2015). The auditory N100 is a negative component occurring 80 to 180 ms after the onset of the stimulus and is maximal at frontal-central sites (Näätänen & Picton, 1987). Its generators appear to be located in the primary and associative auditory cortices, the superior temporal gyrus, and Heschl’s gyrus (Wolpaw & Penry, 1975; Zouridakis et al., 1998). It is thought to reflect the perceptual processing of auditory change detection (Ghani et al., 2020; Näätänen et al., 1978). The P300 is a later positive component unraveling within the 250 – 450 ms time window after the onset of the stimulus and typically increases in magnitude from the frontal to parietal electrode sites (Johnson, 1993). This component is believed to reflect attentional and memory mechanisms (Polich, 2007). The P300 can be divided in two subcomponents, the P3a and the P3b (Snyder & Hillyard, 1976). P3a originates from stimulus-driven frontal attention mechanisms whereas P3b originates from temporal-parietal activity (Polich, 2007). P3a, is thought to reflect an automatic switch of attention from the primary task towards distractive stimuli (Polich, 2003), while P3b seems to reflect the allocation of attentional resources that promote context updating operations and memory processing (Brázdil et al., 2001; Donchin & Coles, 1988; Knight, 1996) and might also index a mechanism linking stimulus identification and response selection (Frühholz et al., 2011; Verleger et al., 2005). According to Squires (1975), the earlier component P3a (latency about 240 ms) is elicited in response to infrequent, unpredictable shifts of either intensity or frequency in a train of tones, whereas the later P3b component (mean latency about 350 ms) occurs only when a subject is actively attending to the tones. In this sense, the P3b is considered to reflect the voluntary detection of a task-relevant stimulus (Picton, 1992; Polich & Criado, 2006), whereas P3a might be more specifically related to deviant auditory non-target events.

While N100 has been found to be modulated by mental load (Dehais et al., 2016), it seems that the literature more frequently find the P300 to be impacted by load or task difficulty (e.g., Kramer et al., 1995; Miller et al., 2011), including in complex applied settings such as aviation (Dehais, et al., 2019; Giraudet et al., 2015; Kramer et al., 1987) or car driving (Van der Heiden et al., 2018; Wester et al., 2008). Regarding the links between sound detection performance and brain response, Dehais, Roy, et al. (2019) have found amplitude reductions of the N100, P3a and P3b for missed target sounds relative to detected ones and Giraudet et al. (2015) revealed that auditory P3b amplitude was negatively correlated with the percentage of missed auditory targets. The latter study also reported a reduction in P3b amplitude when the primary task load increased, while the N100 component remained unaffected, suggesting that load rather affected later cognitive processes than earlier perceptual and/or preattentive processes.

### 1.3. Multiple alarms and auditory load

Exposure to multiple alarms over time can also complicate the identification process (Potnuru et al., 2020). In intense health care settings there can be more than 300 alarms per patient per day, but less than 5% of these alarms require urgent clinical intervention (Association for the Advancement of Medical Instrumentation, 2011). This constant bombardment of alarms leads to ‘alarm fatigue’ (Lansdowne et al., 2016) and also likely create confusion. Mental workload, large number of alarms, and confusion of alarms have been cited as main cause of individual not responding to alarms (Xiao et al., 2003). The auditory environment can vary in complexity, from a quiet environment, with no auditory stimuli other than target ones, to complex environments characterized by numerous sounds conveying independent messages and competing for an operator’s attention (Ho & Spence, 2005). In the latter, the perceptual processing of target stimuli becomes more difficult. The human ear is capable of discriminating pitch differences as small as 0.5% (Wier et al., 1977), and individuals display remarkably accurate pitch memories for environmental sounds, such as the landline dial tone (Smith & Schmuckler, 2008) and isolated tones such as the “bleep” used to censor taboo words in the media (Van Hedger et al., 2015). However, such discrimination becomes almost impossible when numerous alarms sharing acoustic characteristics (Lacherez et al., 2007) are triggered in close succession.

An interesting way to study the detection performance of a tone target among different other auditory stimuli triggered in close succession is also to use the oddball paradigm and its variations (e.g., Ebmeier et al., 1995; Katayama & Polich, 1996). In the 1-tone (or single-tone) paradigm (Cass & Polich, 1997) in which target tones are the only stimuli presented, auditory load is minimal. Participants are required to press a button when the target is presented. In the 2-tone paradigm, two auditory stimuli are played: a standard (non-target) tone occurring frequently, and a target tone occurring less frequently (Squires et al., 1975). Participants are required to respond to the target tone and to ignore the standard tone. This task remains easy since only two audio traces are compared. The 3-tone paradigm includes an additional non-target and rare distractor tone that must be ignored (Courchesne et al., 1984; Katayama & Polich, 1996). While Cass and Polich (1997) found no difference in task performance errors (target misses + false alarms) between the 1-tone paradigm and the 2-tone paradigm, other studies found a few more errors in the 2-tone paradigm than in the 1-tone paradigm (Polich et al., 1994; Polich & Heine, 1996; Polich & Margala, 1997). Katayama & Polich (1996) mention that the 3-tone paradigm can produces a relatively complex stimulus construction compared to the 2-tone situation. They compared detection performance in one-, two-, and three-stimulus auditory paradigms versions. However, they found no difference in target tone detection performance. A possible explanation is that the pitches of the 3 tones were markedly different (500, 1000, or 2000 Hz). Differences in response times (RTs) have been observed, however, with longer RTs in the 2-tone than in the 1-tone paradigm (Polich & Heine, 1996), and in the 3-tone than in the 2-tone paradigm (Ebmeier et al., 1995; Grillon et al., 1990).

Electrophysiological studies using the oddball paradigms and its variants described above found somewhat divergent results. Some studies found no variation in P3b amplitude in response to target tones as the number of irrelevant tones varied (Cass & Polich, 1997; Katayama & Polich, 1996; Polich et al., 1994; Polich & Margala, 1997), while others found a greater P3b responses to the target tone in the 2-tone relative to the 1-tone paradigm (experiment 1 in Cass & Polich, 1997; Polich & Heine, 1996), and in the 3-tone paradigm compared to the 2-tone paradigm (Grillon et al., 1990).

In sum, on balance, the existing literature using oddball paradigms and its variants suggests that auditory load may impact on auditory target detection, with some behavioral and electrophysiological evidence of a reduction of target processing as the number of non-target tones increases.

### 1.4. The present study

As described in the previous sections, both mental and auditory loads can negatively affect an operator’s capacity to detect an alarm. Of importance, however, these two factors have been mainly studied independently and we do not currently well know what their combined impact may have on detection performance, in particular when the different tones played are very close in terms of pitch. The present study combined the manipulation of mental and auditory loads to examine their effects on behavioral performance (target tone performance) and brain activity (event related potentials). Participants were required to perform a dual-task paradigm consisting in an aviation-inspired task and a tone detection paradigm (simulating an auditory alarm detection task). The aviation task − whose difficulty was manipulated to generate different levels of mental load (no load, low load, high load) − was taken from Giraudet et al. (2015). It reproduces a landing situation in which the pilot must decide whether the approach can be continued or should be aborted based on various elements of information displayed in the cockpit (e.g., wind speed, aircraft position, etc.). Concurrently to this aviation task, participants performed a tone detection paradigm whose number of different tones varied (1-. 2-, or 3-tone paradigm, manipulated between-participants) to manipulate auditory load. At the behavioral level, we predicted a greater number of missed target tones as the aviation task’s difficulty (Giraudet, St-Louis, et al., 2015) and the number of different tones increased. At the electrophysiological level, we predicted a decrease in the P3b amplitude in the same conditions. This prediction was based on the hypothesis that (1) an increasing mental load in the aviation task should hinder the redirection of attention from the demanding aviation task towards the target tone in the tone detection paradigm; and that (2) auditory discrimination should be reduced when the number of auditory distractors increases, in particular when 3 different tones are played. Finally, as both the aviation task and the tone detection paradigm tap attentional resources, we predicted additive effects between the mental and auditory loads. More specifically, we predicted a drastic increase in the number of missed target tones and a decrease in P3b amplitude in response to the target tone in condition of high mental load in the 3-tone paradigm.

## MATERIAL & METHODS

### 2.1. Participants

Sixty participants (15 females; *M_age_* = 23.1 years old, *SD* = 3.6) from the University of Toulouse participated in the present study. They were all knowledgeable in the field of aeronautic and had previous experience with flight simulators. All had normal or corrected-to-normal vision and reported no history of neurological or psychiatric disorders. They received no financial compensation for their participation in the study. They were distributed in three different groups, performing the 1-tone, 2-tone, or 3-tone paradigm.

### 2.2. Ethics Statement

The study was conducted in accordance with the Declaration of Helsinki (1973, revised in 1983) and was approved by a national ethics committee (CEEI/IRB00003888). After being informed of their rights, all participants gave their written consent.

### 2.3. Tasks

Two different tasks were used in this study: an aviation task and a tone detection paradigm. Our main measure of interest was the performance in the tone detection paradigm, which was compared across three mental load conditions. In the *no load* condition, participants performed the tone detection paradigm only, thus mental load was minimal. The aviation task was displayed, but participants had to entirely ignore it and to fixate a green cross presented at the center of the screen. In the *low*- and *high-load* conditions, participants performed the tone detection paradigm while concurrently performing the aviation task in which the mental load was low or high, respectively. Participants performed a total of 600 trials, with 200 trials for each difficulty level (no-, low- and high-load). Low and high load conditions were performed first in random order, and the no load condition was always performed after. Within each difficulty condition, trials order was counterbalanced. The aviation task was displayed on a 24’ computer screen and the tones were played via speakers placed on both sides of the screen. Both tasks were programmed in Matlab, using the Psychophysics Toolbox extensions (Brainard, 1997; Kleiner et al., 2007; Pelli & Vision, 1997). Mental load condition was a within-participant factor and the tone detection paradigm condition (i.e., number of tones) a between-participants factor (i.e., three different groups).

#### 2.3.1. Aviation task

This task was designed to reproduce a decision-making task performed by pilots during the landing phase, but it can be performed by non-pilot subjects. The participants were presented with short video clips displaying a Primary Flight Display (PFD), an instrument located in front of pilots in aircraft cockpitsn and where the main flight parameters are displayed. Compared to an actual cockpit display, the PFD was simplified (various parameters were omitted). The information available on this simplified PFD included the aircraft’s vertical and horizontal position relative to the landing field (represented by two moving cursors; one on a vertical axis, on the right of the artificial horizon, and the other on a horizontal axis, below the artificial horizon, see Figure 1). In addition, three indicators were displayed on the top left of the screen: the heading (“Cap”, optimal value = “180”), wind speed (“Vent”, optimal value = “0.00”), and magnetic deviation (displayed below the heading, optimal value = “0”). The magnetic deviation is particular, its value must be subtracted (when negative) or added (when positive) to the heading value. The three indicators were static during the video, only the two cursors were moving.

**Figure 1.**
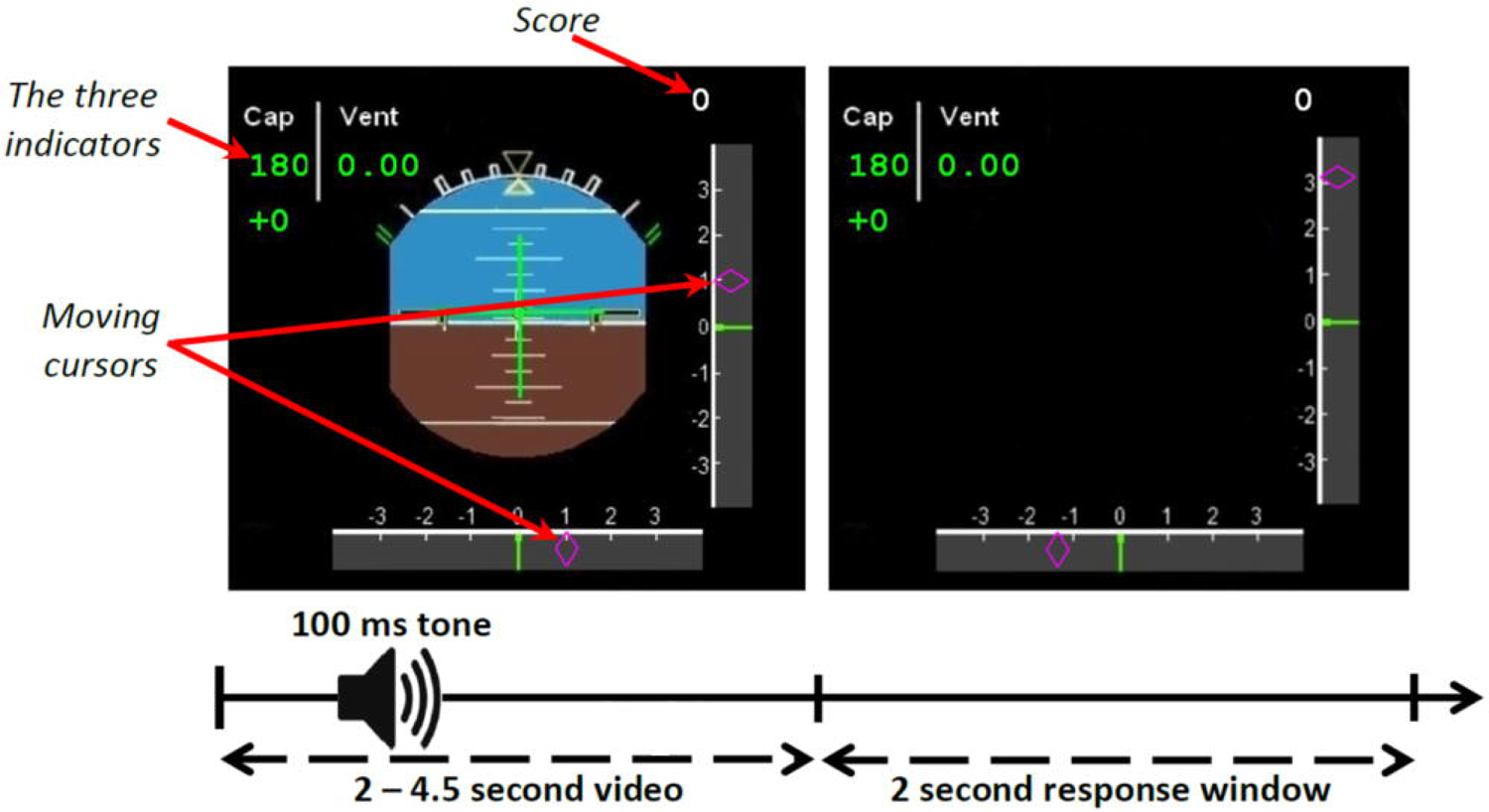
Illustration of a trial in the low load condition. A 2 to 4.5-second video was displayed during which a tone was played, followed by a 2-second response window during which the participant responded to the landing task and then pressed again a button if he/she detected a target tone.

Each trial consisted of a 2- to 4.5-second video clip, followed by a 2-second response time window during which participants had to respond by pressing a corresponding button (see Fig. 1): right (accept landing) or left (abort landing). At the onset of the 2-second response window, the moving cursors were frozen, and the artificial horizon within the PFD disappeared, thereby prompting participants to respond. Participants were asked to respond based on the information displayed during this response window (see below).

In low mental load trials, participants were asked to make their decision based solely on the final position of the two cursors marking the aircraft vertical and horizontal position. The three indicators located on the upper left corner, which participants were instructed to ignore, appeared in green and their values deviated by less than 5 units from their optimal ones. The landing was considered possible when the deviation of each cursor was in a [-2, +2] interval on the arbitrary scale (see Figure 1). In the high load trials, the cursors moved two times faster than in the low load trials, and the three indicator values appeared in red, which indicated a degradation of the aircraft situation. In this case, participants had to make their decisions based on both the cursors and the indicators, checking whether heading and wind values deviated by more than 5 units from their optimal value (acceptable range for wind: 0-5; acceptable range for wind: 175-185). An illustration of the low- and high-load trials as well as the rules to be followed by participants to make a decision are presented in Figure 2.

**Figure 2.**
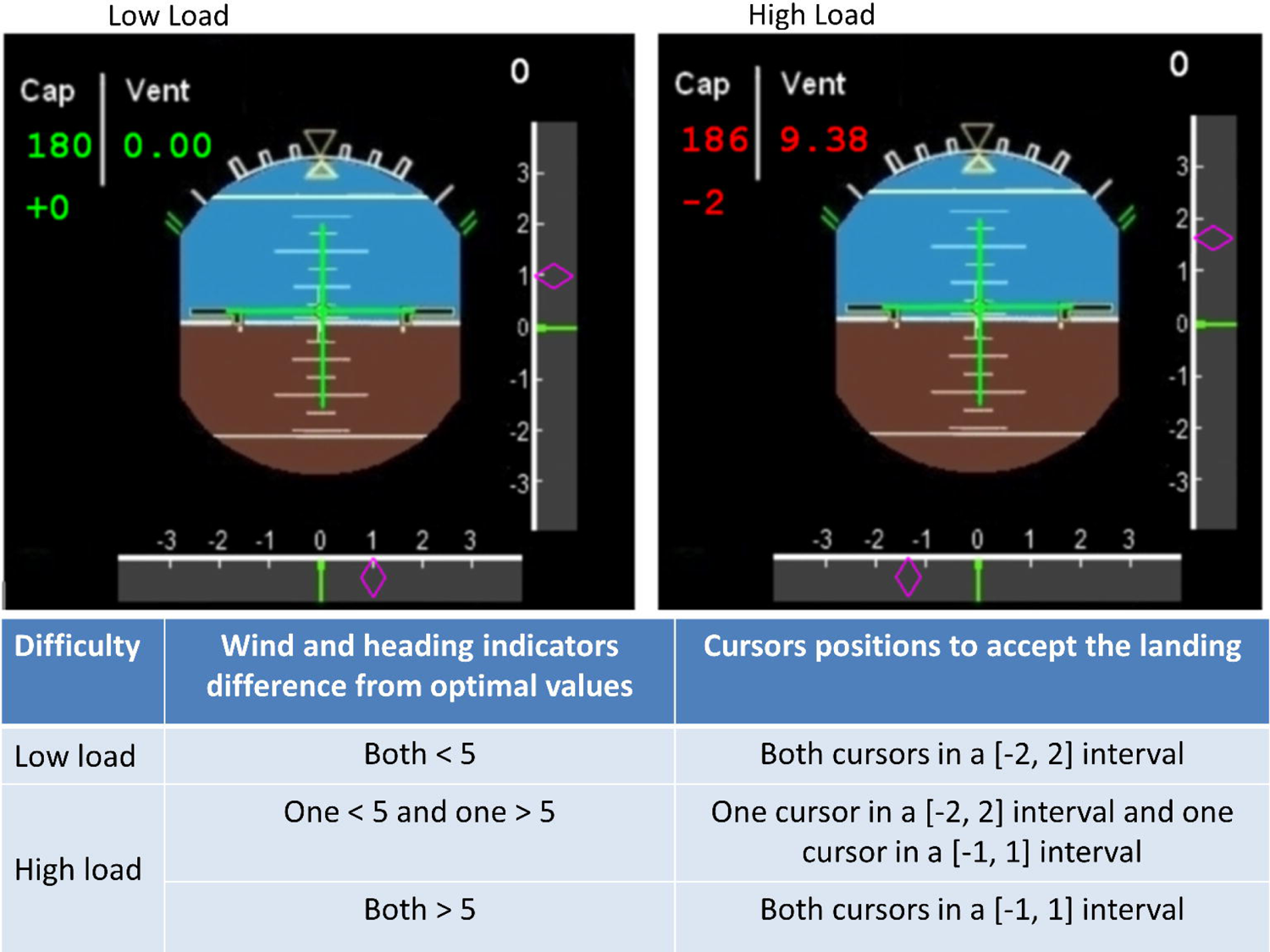
Top: Still capture of examples of videos used in the low- and high-load conditions (left and right pictures, respectively). Bottom: table indicating the criteria to base landing decision on. In the low-load example (top left), indicator values are presented in green, indicating that they are optimal. Landing is acceptable, for the cursors marking the aircraft’s horizontal and vertical positions are both within the [-2, +2] range. In the high-load example (top right), the wind value is 9.38, thereby deviating by more than 5 units from the optimal 0 value; the heading value is 184 (i.e., 186 – 2 magnetic deviation units = 184) and thus deviates by less than 5 units from the optimal 180 value. In this situation, with one indicator outside the acceptable range and the other within it, landing should be carried out only if one of the two cursors falls within [-2, +2] range while the other falls within a more conservative [-1, +1] range. The vertical position is deviating by less than +2 units but the horizontal position is deviating by more than −1 unit, consequently, the landing conditions are not met and landing should be aborted. In the event that both heading and wind values are outside the acceptable range, tolerance on cursor deviation is even lower and landing should only accepted if both cursors fall within the [1,+1] range.

To increase motivation and engagement in the task, the participants’ score (number of correct decisions) was displayed on the upper right corner of the PFD (see Figure 1). For each correct landing response, the score increased by 1 point. When participants missed a response, or made an incorrect response, this score remained unchanged.

#### 2.3.2. 1-, 2-, and 3-tone detection paradigms

Depending on the group they were assigned to, participants performed the 1-, 2, or 3-tone version of the tone detection paradigm. During each 2 to 4.5-seconds video, one 50 dB (SPL) tone lasting 100 ms was randomly played somewhere between 500 ms after the onset of the video and 500 ms before the end of the video, see Figure 1. Note that only one tone type was presented during each video, and the term audio load is used in this paper to describe the difficulty to identify a target tone while other very similar tones can be also played. In each trial, the auditory stimulus could be a standard tone (sound frequency = 1900 Hz), a target tone (sound frequency = 1970 Hz), or a deviant tone (sound frequency = 2040 Hz), depending on the tone detection paradigm performed by the participant. The pitch of the tones was inspired by a study of Kolev et al. (1997). In the 1-tone paradigm, only targets tones were played (n = 60). In the 2-tone paradigm, the tones were targets (n = 60; probability = .30) or standards (n = 140; probability = .70). In the 3-tone paradigm, the tones were targets (n = 40; probability = .20), deviants (n = 20; probability = .10), or standards (n = 140; probability = .70). The number of target tones was the same across the three levels of mental load (e.g., a participant that performed the 1-tone paradigm was administered with 60 target tones in the no mental load condition, 60 target tones in the low mental load condition, and 60 target tones in the high mental load condition). The pitch of the target tone was maintained (i.e., 1970 Hz) across the three tone detection paradigms to avoid introducing a bias that could have influenced ERP responses. We also maintained the same probability (i.e., .70) for the standard tones across the 2- and the 3-tone paradigms. During the response window, participants were instructed to respond to the landing task first (except in the no load condition during which the aviation task was presented but had to be ignored), and then to report any potential target tones by pressing a push button. They were asked to ignore the standard tones (in the 2- and 3-tone paradigms) and the deviant tones (in the 3-tone paradigm). To minimize motion artifacts on the EEG signal, we separated the video clip period, during which tones were played and ERP analysis performed, from the 2-s response time window. The performance to the alarm detection task was measured with the percentage of miss and false alarms (other performance measures, including the d’, are available in supplementary material).

### 2.4. Procedure

Participants were seated in a comfortable armchair in a sound-dampened room. They completed the consent form and were instructed to keep their forearms stable on the chair’s arms, with their two hands resting on the response box (Cedrus RB-740). Participants were introduced with the tone(s) corresponding to their tone detection paradigm, and we ensured that they were able to recognize it/them. Then, participants underwent a training consisting of 10 trials of the no mental load, low mental load, and high mental load conditions. They performed the tone-detection task as during the actual experiment. Following this training, participants were fitted with the EEG electrode cap as well as the Electro-OculoGraphic (EOG) electrodes (see description below). They then performed the low mental load and high mental load conditions (random order, 40 minutes), during which they had to perform the aviation task and the tone detection paradigm concurrently. Participants were instructed to give equal importance to the aviation task and tone detection paradigm. Next, they performed the no mental load condition (20 min) in which they performed the tone detection paradigm only (while ignoring the aviation task). The whole procedure lasted approximately 100 min, including 20 min to install the EEG (see Figure 3).

**Figure 3.**
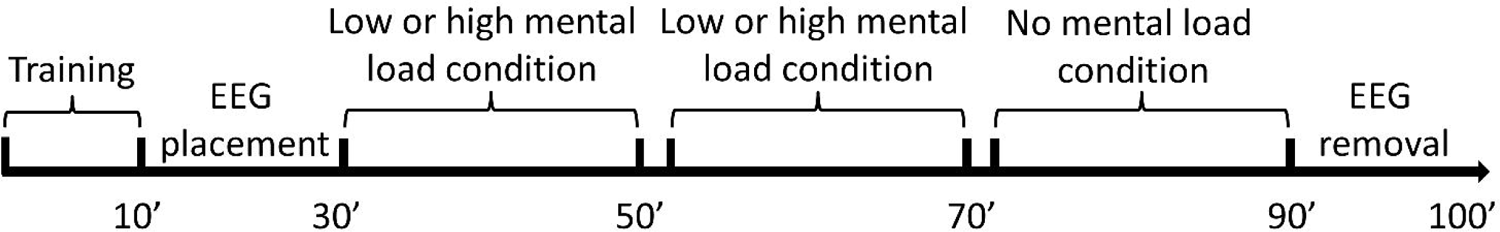
The procedure timeline.

**Figure 4.**
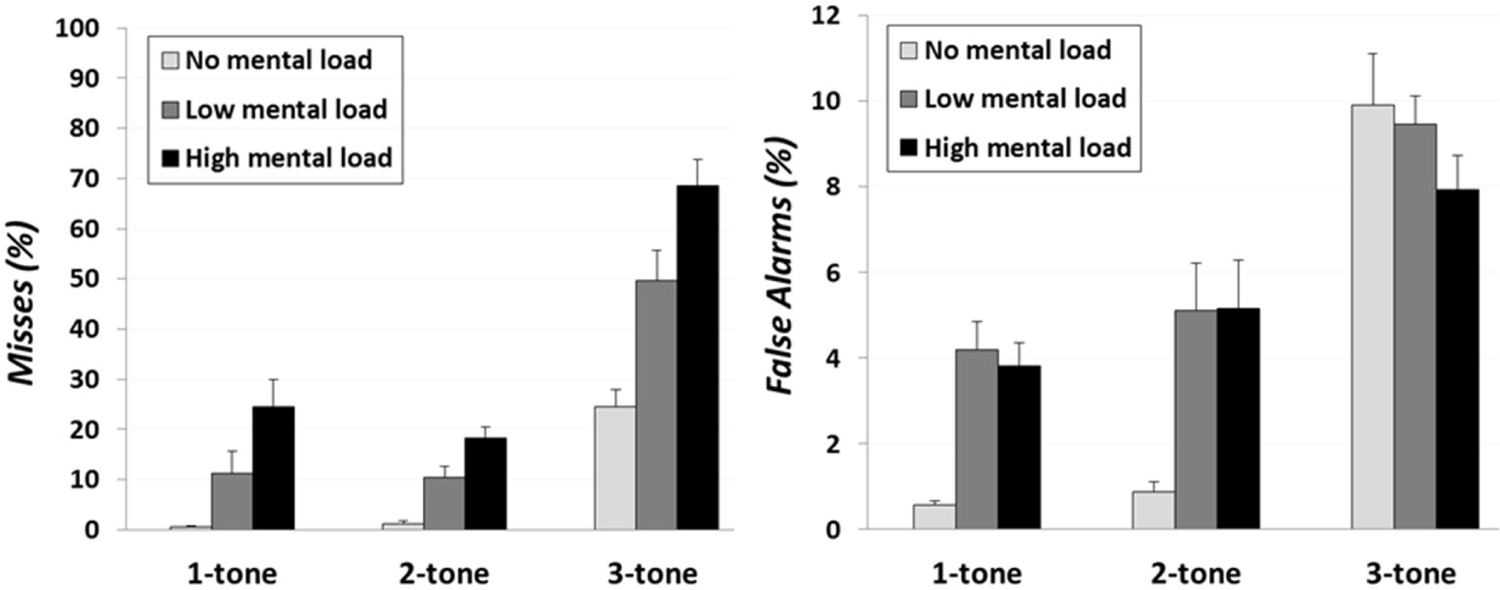
Detection of the target tone in the no load (light grey), low load (medium grey) and high load (black) conditions for the 1-tone, 2-tone and 3-tone paradigms. Left: Misses. Right: False Alarms. Error bars are s.e.m.

### 2.5. Data Acquisition

#### 2.5.1. Electroencephalographical recordings

Electroencephalographical (EEG) data were recorded with a BioSemi ActiveTwo system (http://www.biosemi.com) from 64 Ag/AgCl active electrodes (Fp1, AF7, AF3, F1, F3, F5, F7, FT7, FC5, FC3, FC1, C1, C3, C5, T7, TP7, CP5, CP3, CP1, P1, P3, P5, P7, P9, PO7, PO3, O1, Iz, Oz, POz, Pz, CPz, Fpz, Fp2, AF8, AF4, AFz, Fz, F2, F4, F6, F8, FT8, FC6, FC4, FC2, FCz, Cz, C2, C4, C6, T8, TP8, CP6, CP4, CP2, P2, P4, P6, P8, P10, PO8, PO4, and O2) mounted on a cap and placed on the scalp according to the international 10-20 system, plus two sites below each eye to monitor eye movements. Two additional electrodes placed close to Cz − the common mode sense (CMS) active electrode and the driven right leg passive electrode − were used to drive the participants’ average potential as close as possible to the AD-box reference potential (Metting Van Rijn et al., 1991). Electrode impedance was kept below 5 kΩ for scalp electrodes and below 10 kΩ for the four eye channels. Skin–electrode contact, obtained using conductive gel, was monitored, keeping voltage offset from the CMS below 25 mV for each measurement site. All the signals were DC-amplified and digitized continuously at a sampling rate of 512 Hz, using an anti-aliasing filter (fifth-order sinc filter) with a 3-dB point at 104 Hz. No high-pass filtering was applied online. Data were analyzed with the EEGLAB toolbox (Delorme & Makeig, 2004). EEG data were re-referenced offline to the average activity of the two mastoids and bandpass filtered (0.1 - 40 Hz, 12 dB/octave), given that the low-pass filter was not effective in completely removing the 50 Hz artifact for some participants. An independent component analysis was performed to isolate eye-blinks and movements related artifacts that were removed from the signal. A visual inspection of the data was done to reject residual artifacts intervals. Data were segmented into epochs from − 200 to 800 ms that were time-locked to the onset of the target tone. A 200 ms pre-stimulus baseline was used in all analyses. Data with excessive blinks were adaptively corrected using independent component analysis. Segments including artifacts (e.g., excessive muscle activity) were eliminated offline before data averaging. A total of 15% of data were lost due to artifacts. Five participants were removed from the analysis due to the poor quality of the EEG signal. In the end, 19 participants, 18 participants and 18 participants were respectively tested in the 1-tone, 2-tone and 3-tone variants of the tone detection paradigm. In order to investigate different processing stages of the target sounds, we focused on the N100 and the P3b responses.

## 3. RESULTS

### 3.1. Aviation task

A 2 (mental Load: low load, high load) x 3 (auditory load: 1, 2, 3) mixed ANOVAs revealed a significant main effect of the mental load on the number of errors committed in the aviation task, *F*(1, 52) = 20.43, *p* < .001, 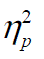= .27, with more errors under the high mental load than under the low mental load. The main effect of the auditory load was not significant *F*(1, 52) = 2.02, *p* = .141, 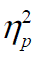= .07, but it interacted with mental load, *F*(2, 52) = 24.23, *p* < .001,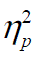 = .46. Whereas decision errors increased as mental load increased in the 1- and 2-tone paradigm (LSD: *p* < 0.001 for both comparisons), the opposite pattern was observed in the 3-tone paradigm (LSD: *p* = 0.004).

### 3.2. Target tone detection performance

All detection performance values are in supplementary material S1. The percentage of errors, misses (number of misses/(number of hits + number of misses)) and false alarms (number of false alarms/(number of false alarms + number of correct rejections), were analyzed using a 3 (Mental Load (no load, low load, high load) × 3 (Auditory load: 1, 2, 3) mixed ANOVAs – with mental load as a within-participant factor and the auditory load as a between-participants factor. All post-hoc analyses were performed using LSD.

*Miss rate.* We found a significant main effect of mental load, *F*(2, 104) = 69.95, *p* < .001, 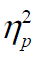 = .57, showing that a higher percentage of misses in the high mental load vs low mental load condition and in the low mental load vs no mental load condition (both *p_s_* < .001) (see Figure 5, right). The main effect of the auditory load was significant, *F*(2, 52) = 46.98, *p* < .001,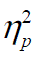 = .64, with a higher miss rate in the 3-tone paradigm than in both the 1-tone and 2-tone paradigms (both *p_s_* < .001). Finally, the interaction term was also significant, *F*(4, 104) = 4.62, *p* = .001, 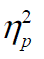 = .15, showing more important effects of increased mental load on the miss rate in the 3-tone paradigm than in both the 1-tone and the 2-tone paradigms.

**Figure 5.**
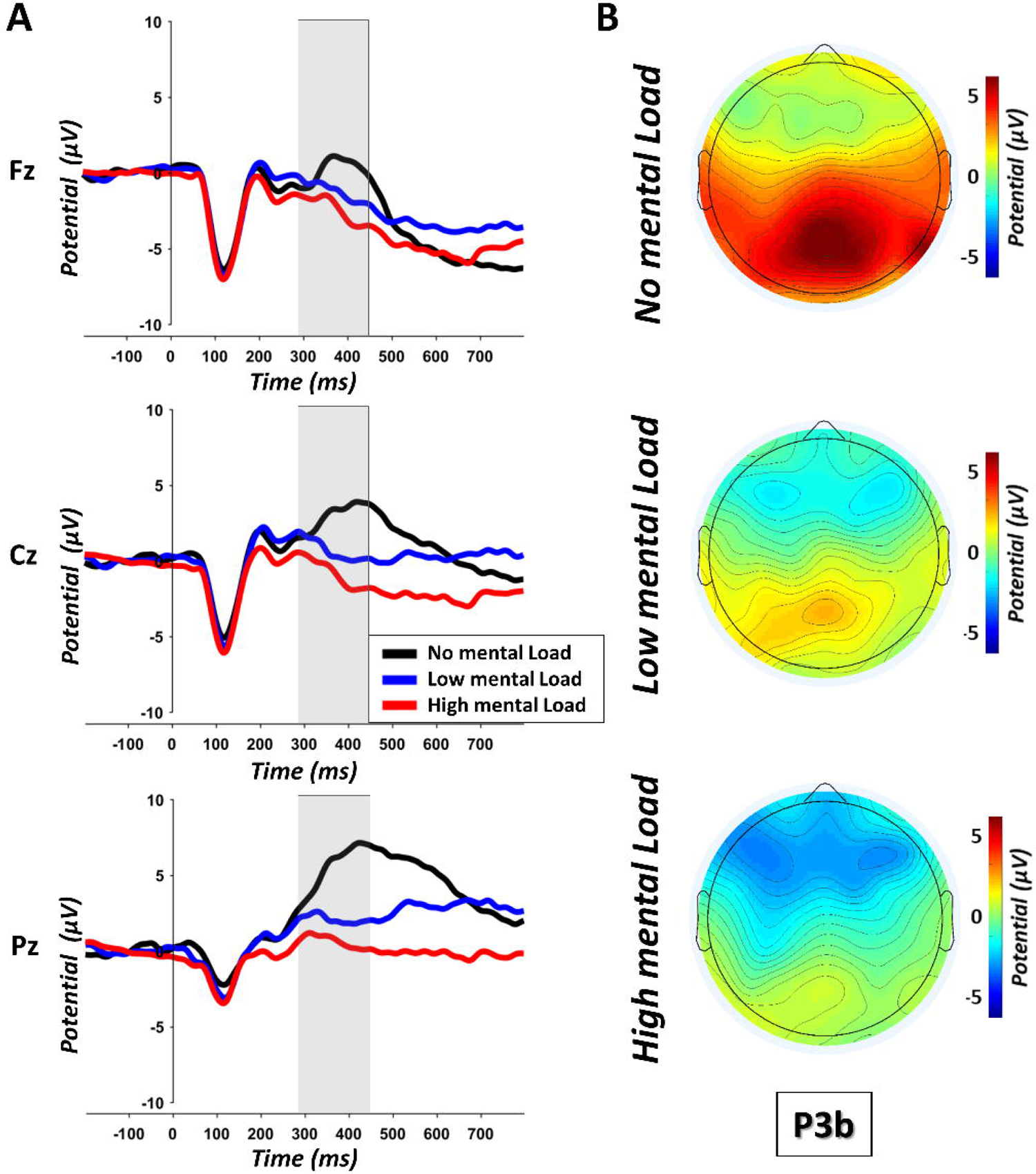
(A) Grand average ERP waveforms at Fz, Cz and Pz observed in response to the target tones in the no (black line), low (blue line), and high mental load conditions (red line). (B) Scalp maps illustrating the P3b mean amplitude in the no (up), low (middle) and high mental conditions (down). A bar chart of the average P3b values at Fz, Cz and Pz during each tone detection paradigm is in supplementary material S3.

*False alarm rate.* A significant main effect of mental load was found, *F*(2, 104) = 13.79, *p* < .001, 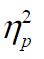 = .21, with more false alarms in low and high mental load conditions than in the no load mental condition (both *p_s_* < .001; see Figure 5, bottom left). The main effect of the auditory load was significant, *F*(2, 52) = 29.01, *p* < .001, 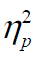 = .53, with a markedly higher number of false alarms in the 3-tone paradigm than in the 1-tone and 2-tone paradigms (both *p_s_* < .001). The interaction term was significant, *F*(4, 104) = 8.47, *p* < .001, 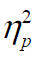 = .24, while the false alarm rate increased between the no mental load and the low/high mental load conditions in the 1-tone and 2-tone paradigms (all *p_s_* < .001), a rather opposite pattern was found in the 3-tone paradigm in which the false alarm rate decreased between the no mental load and high mental load conditions (*p* = .021).

### 3.3. Electrophysiological results

N100 and P3b values are in supplementary material S2. The N100 component was assessed with reference to the peak amplitude within the 80 – 150 ms time window at the Fz, Cz and Pz electrodes (e.g., Valéry et al., 2017). The P3b was assessed with respect to the mean amplitude within the 300 – 450 ms time window at Fz, Cz and Pz electrodes (Giraudet et al., 2016). The amplitudes of the N100 and P3b components were analyzed at each of these electrode sites using a 3 (Electrode: Fz, Cz, Pz) × 3 (Mental Load: no load, low load, high load) × 3 (Auditory Load: 1, 2, 3) mixed ANOVAs − with electrode and mental load as within-participant factors and the auditory load as a between-participants factor. All post-hoc analyses were performed using LSD.

#### 3.3.1. N100 component

We found a significant main effect of electrode, *F*(2, 104) = 91.22, *p* < .001, 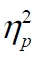 = .64, with greater N100 amplitudes in response to the target sound at Fz than at Cz (*p* < .001) and at Cz than at Pz (*p* < .001). The main effects of mental and auditory loads were not significant *F*(2, 104) = 0.29, *p* = .743, 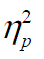 = .01, and *F*(2, 104) = 0.83, *p* = .438, 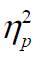 = .03, respectively (see Figure 6 and 7). However, we found a significant Electrode x Auditory Load interaction, *F* (4, 104) = 7.08, *p* < .001, 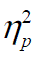 = .21, with reduced N100 amplitudes only at Fz in the 1-tone paradigm relative to the 2-tone and 3-tone paradigms (both *p_s_* < .050), see Figure 7. The other interaction terms were not significant: Mental Load x Auditory Load *F*(4, 104) = 0.86, *p* = .489, 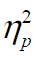 = .03, Mental Load x Electrode, *F*(4, 208) = 0.19, *p* = .943, 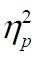 = .00, and Mental Load x Auditory Load x Electrode *F*(8, 208) = 0.71, *p* = .676, 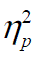 = .03.

**Figure 6.**
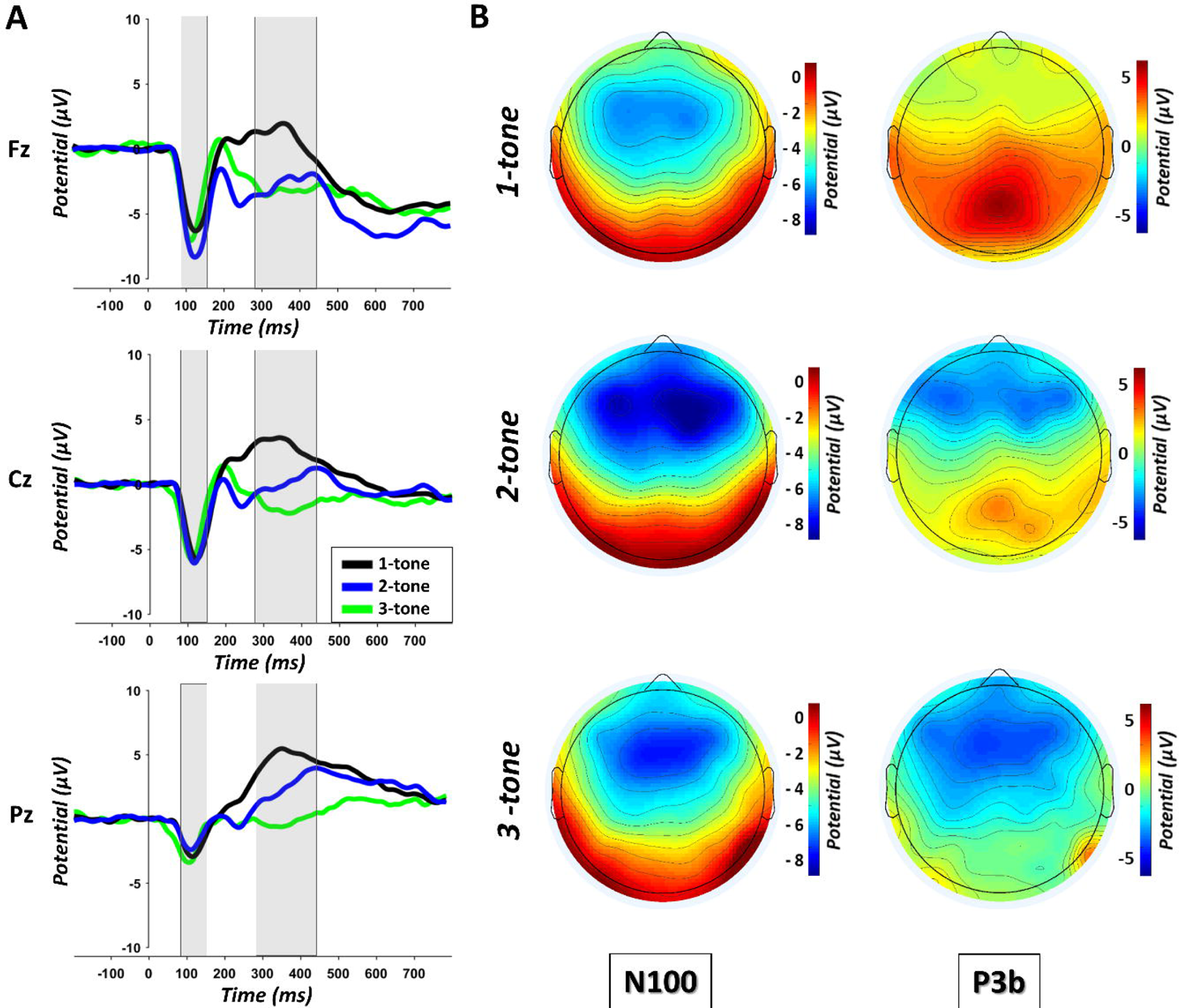
(A) Grand average ERP waveforms at Fz, Cz and Pz observed in response to the target tone in the 1-tone (black line), 2-tone (blue line) and 3-tone paradigms (green line). (B) Scalp maps illustrating the N100 peak amplitude and the P3b mean amplitude in the 1-tone (up), 2-tone (middle), and 3-tone paradigms (down). A bar chart of the average P3b values at Fz, Cz and Pz during each tone detection paradigm is in supplementary material S4.

**Figure 7.**
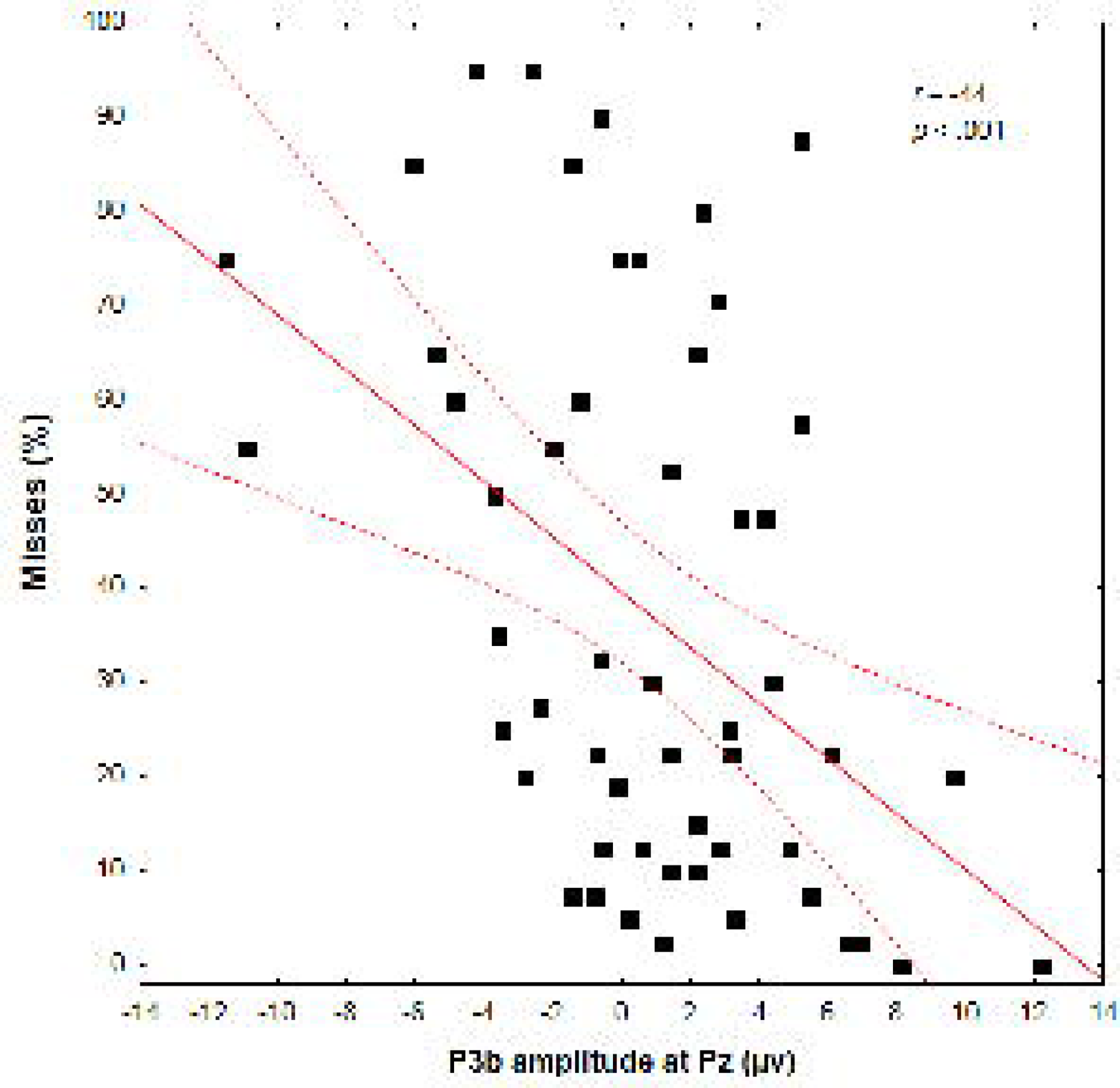
Correlation between the miss rate (missed target tones) and the mean P3b amplitude for all tone detection paradigms. N = 54.

#### 3.3.2. P3b component

The analysis of the P3b amplitude revealed a significant main effect of electrode *F*(2, 104) = 100.55, *p* < .001, 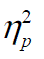 = .66, with a greater P3b amplitude in response to the target sound at Pz than at Cz and at Cz than at Fz (both *p_s_* < .001). The main effect of mental load was significant too, *F*(2, 104) = 20.21, *p* < .001, 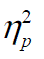 = .28, with lower P3b amplitude under high than low mental load (*p* = 0.039) and under low than no mental load (*p* < .001, see Figure 6). The P3b amplitude also varied with the auditory load, *F* (2, 52) = 26.95, *p* < .001, 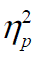 = .51, with a significantly greater P3b amplitude in the 1-tone than in the 2-tone paradigm and in the 2-tone than in the 3-tone paradigm (*p* < .001 and *p* = .006, respectively, see Figure 7). The Electrode x Auditory Load interaction was significant, *F*(4, 104) = 3.03, *p* = .021, 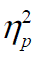 = .10. The P3b was significantly greater in the 1-tone than in the 2-tone paradigm, and in the 2-tone than in the 3-tone paradigm on all electrodes, this effect being larger at Pz, see Figure 7. The Electrode x Mental Load interaction was significant too, *F* (4, 208) = 8.99, *p* < .001, 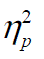 = .15. Reduced P3b amplitude was found under high than low mental load, and under low than no mental load, for all electrodes, these effects being larger at Pz. Finally, neither the Mental Load x Auditory Load interaction nor the triple interaction were significant, *F* (4, 104) = 0.64, *p* = .631, 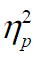 = .02, and *F* (8, 208) = 0.96, *p* = .462, 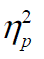 = .04, respectively.

#### 3.3.3. Correlation between the percentage of hits and P3b amplitude

We performed two Pearson correlations to examine the relationship between individual miss rate and P3b amplitude (Pz electrode) elicited by the target tones in the low and high mental load conditions. This analysis included the three groups of participants (i.e., the three tone detection paradigm variants). We found a significant negative correlation between miss rate and P3b amplitude both in low (*r* = − .29, *p* = .036) and high load conditions (*r* = − .44, *p* < .001). In other words, the more the participants missed target tones, the lower was their mean P3b amplitude at the Pz electrode, see Figure 9.

## 4. DISCUSSION

The aim of the present study was to investigate the impact of both mental and auditory loads on alarm responsiveness using behavioral and electrophysiological measures. This was achieved using a simplified aviation task (Giraudet et al., 2015) with a concomitant tone detection paradigm (simulating an alarm detection task). Mental load varied a function of the aviation task difficulty and the auditory load was manipulated with the number of different tones possibly played in the tone detection paradigm. Overall, our results showed that the detection of auditory target stimuli is hindered by both mental and auditory loads, and that the interaction of these factors yields the greater reduction in detection performance. Our electrophysiological results showed that, by and large, the N100 response was immune to the impact of these two factors. In contrast, the amplitude of the P3b reduced as both loads and increased, these effects being additive. As a reminder, the P300 subcomponent elicited in this study was the P3b since it is considered to be elicited by the voluntary detection of a task-relevant stimulus (Picton, 1992; Polich & Criado, 2006).

### 4.1. The impact of mental and auditory loads on target tone detection performance

In line with our predictions, increasing the aviation task load negatively impacted participants’ performance to the tone detection task. Interestingly, increased miss rate was concomitant with an increased false alarm rate, thus this lower detection performance was not due to a lack of time to report the target tones or to a change toward a more conservative response criterion. In line with Giraudet et al. (2016), we found no modulation of the N100 amplitude to the target tones by mental load, suggesting that early perceptual processes were not significantly impacted by this factor. In contrast, the P3b response decreased in amplitude as mental load increased, especially in the parietal region (Pz electrode). At least two explanations can account for the tight relationship between increased mental load and reduction of both tone detection performance and P3b amplitude. First, given the parietal contribution to the P3b generation (Polich, 2007), and the role of this cerebral structure in the control of voluntary attentional orienting (Corbetta et al., 2000), the modulation of the P3b amplitude by mental load probably relate to the increasing difficulty to voluntary orient attention towards the target tones when the aviation task demand on attentional resources increased, and thus when attentional resources became scarce (Corbetta et al., 2000), assuming that some limited attentional resources are shared between the aviation task and the tone detection paradigm. Second, an increased mental load might also affect both the storage of the target tone representation in working memory and the selection of the associated response (Frühholz et al., 2011; Verleger et al., 2005). The correlation analysis revealed that increased alarm miss rate was associated with lower P3b amplitude. This result supports the idea that the P3b amplitude is a good indicator of the success to detect an auditory target. It is consistent with previous studies reporting lower P300 amplitudes for missed than for detected alarms (Dehais, Roy, et al., 2019), or significant correlations between the miss target tone rate and P300 amplitude (Giraudet, St-Louis, et al., 2015).

A second key finding in our study is the decrease in auditory detection performance as the auditory load increased, in particular when 3 different tones were played. While, in the no mental load condition, the miss rate was very low in the 1-tone (0.53%) and 2-tone (1.11%) paradigms, in line with the error rate observed by Polich & Heine (1996) in similar conditions (an experiment without secondary task, as in our no mental load condition), the miss rate increased drastically (24.44%) in the 3-tone paradigm. This effect was accentuated as load increased, yielding to the higher miss rate in the 3-tone paradigm under the high mental load condition (68.64%). In this situation, participants struggled to properly detect and discriminate the target tones from the two irrelevant ones. These results contrasted with the study from Katayama & Polich (1996) in which the miss rate did not decline in a 3-tone paradigm. It is very likely that the main reason for this difference is that the pitch of the three tones were markedly different in their study (500, 1000, or 2000 Hz) while it was very close in our (1900, 1970, 2040 Hz), creating much more difficulty to discriminate them in the 3-tone paradigm.

This impact of auditory load on detection performance was marginally reflected by the N100 and markedly by the P3b responses. Lower N100 amplitudes (less negative) were found in response to the target tone in the 1-tone than in both the 2- and 3-tone paradigms. This result is complex to interpret but we might argue that the N100 was influenced by the MMN, which is elicited by when repetitive standard auditory stimuli are occasionally interspersed with deviant stimuli (May & Tiitinen, 2004). Supporting this idea, some authors hypothesize that N100 and MMN are produced by common neural mechanisms (May & Tiitinen, 2004). In the 1-tone paradigm, only one tone was played, which could explain the reduced N100. Importantly, the amplitude of the P3b response decreased as the auditory load increased, suggesting a reduced identification of the target tone as the number of additional tones increased. The lower target detection performance found in the 3-tone paradigm may have different explanation than the lower target detection performance due to increase mental load. We used brief sinewave tones that were previously unknown to the participants. Without knowledge of these tones in long term memory, participants must maintained the three auditory traces in auditory working memory (Van Hedger et al., 2018) to compare them with the currently played tone, which was probably very difficult since the tones were very similar in terms of pitch. It has been suggested that perceptual similarity between items result in a large amount of overlapping information, which can lead to greater comparison errors between items encoded into working and the test items presented at retrieval (Jackson et al., 2015). This may have contributed to a weaker memory representation of the target tone in working memory and consequently a lower ability to engage an appropriate response (Frühholz et al., 2011; Verleger et al., 2005).

### 4.2. Practical and neuroergonomics applications

Our results provide empirical evidence of the increasing difficulty for operators to detect alarms when both mental and auditory loads increase. Importantly, aspects of our results indicate that these factors are additive and amplify the risk of missed alerts. This may provide useful information to help improving the design of alarm systems in a wide range of safety-critical domains, such as transportation (e.g., semi-automated vehicles, aviation), medical care or nuclear powerplants. While it seems difficult to predict individual susceptibility to miss alerts, for example, individual working memory capacity seems not to be a relevant predictor (Kreitz et al., 2016), one possible measure to limit alarm may consist in monitoring and balancing the operators’ mental load in the working environment to spare attentional resources and ensure the appropriate detection and processing of unexpected auditory alarms (Giraudet, St-Louis, et al., 2015). The real-time monitoring of an operator’s mental load may possibly be implemented using a combination of performance monitoring, a physical assessment of the amount of stimulation and task difficulty faced by operators, and the measurement of key physiological indicators such as in this study. An optimum level of mental load and should be sought, so to avoid overload or under-stimulation, which can increase the risk of operators drifting “out of the loop” (Gouraud et al., 2017). Exposing regularly operators with the different alarms may also help maintaining the auditory traces in long term memory (Van Hedger et al., 2018), which should facilitate their recognition especially when some alarms are very close in terms of acoustic properties. Another recommendation would be to reduce the acoustic traffic jam in saturated settings such as operating rooms or cockpits, by limiting the number and the variety of auditory alarms that can be issued during specific time periods (Edworthy, 2013; Konkani et al., 2012). Recent results showed that our capacity to process auditory alarms depends in part on the relevance of the auditory stream (Scheer et al., 2018; Tellinghuisen et al., 2016), individuals are less sensitive to auditory stimuli when instructed to disregard auditory information as opposed to monitor it. Scheer et al., (2018) suggest that requiring pilots to perform simple and frequent tasks in the auditory modality could heighten their awareness of the auditory environment and prevent the occurrence of inattentional deafness. However, while this would serve the purpose of increasing their detection at some point, it also means a higher risk of saturating the auditory scene or possible interference with the processing of other auditory signals. Our study critically shows that brief pure tones can largely fail to attract operators’ attention or can be confused, under demanding situation. Thus, a solution may also be to improve the design of alarm systems. According to Foley et al. (2020) “re-orchestring” standardized alarms with more complex sounds may be a desirable way to improve alarm efficiency and avoid confusion (Lacherez et al., 2007). Also, verbal (speech) warning signals can communicate more precisely some information than non-speech warnings (e.g., pure tones) about an upcoming event. Interestingly, some particular words might be efficient to break through the attentional barrier and capture the attention of an operator. For example, Moray’s (1959) well-known study of the “*cocktail party phenomenon*” showed that participants can notice their name embedded in an ignored auditory channel. Later, Wood and Cowan (1995) showed that participants who recalled hearing their name in a to-be-ignored auditory channel (which was the case in 34.6% of the participants) were able to monitor this irrelevant channel during a short period of time afterward. Calling the name of a pilot might be an easy and efficient solution to improve alarm detection in critical situations. Other authors recommend the use of natural auditory indicators (e.g., using the sound of a heartbeat to notify to medical doctors that a patient’s cardiac activity is abnormal) to maintain a correspondence between the acoustic property of the auditory natural indicator and the acoustic property of the message (Fitch & Kramer, 1994; Stevens et al., 2009). Some evidence suggests that such natural auditory stimuli can reduce reaction times in operators compared to tonal or speech warnings (Graham, 1999). Using auditory natural indicators in the cockpit when pilots are under high levels of load might be a suitable solution to prevent inattentional deafness. On a more general note, as the brief pure tones used in the present study were not really representative of real alarms (Ho & Spence, 2005), it would be useful in future studies to include more complex and ecologically-valid sounds (Winkler, 2003). Finally, evaluating and validating alarm systems using brain electrophysiological measurements would be a promising way to enhance alarms and prevent missed alerts. It may be useful for industry stakeholders and practitioners to set up alarm evaluation platforms and perform more systematically empirical experiments similar to the present one to determine the efficiency of specific alarms using behavioral and cerebral data (in particular P3b component). Light and portable EEG devices are now relatively cheap and easy to deploy, and such assessments may help increase the likelihood that alarms are detected by operators even in situations of high mental load. It may also help assessing whether alarms sufficiently distinct from one another to reduce the risk of auditory confusion (Lacherez et al., 2007).

### 4.3. Conclusion

The results of the present study demonstrate that operators’ mental and auditory loads alter the ability to respond to auditory alarms, most likely because these factors affect the attention orientation mechanism, the identification process, the storage of the target sound representation in working memory, and the subsequent selection of the associated response, as suggested by reductions in P3b amplitude. We also found that P3b amplitude was correlated with individual alarm miss rate. Taken together, our behavioral and ERP results support the idea that the inability of pilots to react to some auditory alarms in the cockpit during critical flight phases is also due to cognitive and attentional limitations, contrary to the well-known cry-wolf effect, during which operators or pilots deliberately ignore alarm considered as unreliable due to a high rate of false positives (Bliss et al., 1995, 1999).

Measuring the P3b response may constitute an efficient tool to evaluate the ability of an individual to process auditory alarms in high load contexts. Also, measuring P3b may also help to evaluate alarm designs, and objectively identify the ones that can be efficiently perceived despite such high load contexts. Our data also confirmed that studying alarm responsiveness can be performed with a 1-tone detection paradigm, and the latter is probably more suited to study inattentional deafness. Indeed the classical 2-tone detection paradigm (i.e. oddball tasks), sometimes used to study this inattentional deafness, may be more suited to study the “change deafness” phenomenon (Vitevitch, 2003), since two auditory traces are compared, which is quite different than the simple detection of a single stimulus as usually found in the inattentional deafness literature (Macdonald & Lavie, 2011).

Numerous domains could benefit from a better consideration of the limitations of the human brain in terms of responsiveness to auditory alarms, including environments where human and artificial systems share control. For example, autonomous vehicles can emit alarms to initiate a take-over of control by the human driver in case of emergency. However, if operators are engaged in another task, they might not be able to detect an alarm and their responsiveness may be reduced (Janssen et al., 2019; Van der Heiden et al., 2018).

## Data availability statement

The datasets generated for this study are available on request to the corresponding authors.

## Authorship contribution statement

Mickaël Causse: Conceptualization, Methodology, Software, Data curation, Formal analysis, Writing – original draft, Supervision, Resources. Fabrice B. R. Parmentier: Writing – review & editing. Damien Mouratille: Software, Data collection, Data curation, Formal analysis. Dorothée Thibaut: Data collection, Formal analysis. Marie Kisselenko: Data collection, Formal analysis. Eve Fabre: Methodology, Data collection, Data curation, Formal analysis, Writing – original draft, Supervision.

## Acknowledgments

The authors would like to thank Johanna Clerc for her help with data collection.

---

**Tone detection performance**
| **Correct Hit Rate (%)** | 1-Tone paradigm |  | 2-Tone paradigm |  | 3-Tone paradigm |  |
| --- | --- | --- | --- | --- | --- | --- |
|  | *M* | *SD* | *M* | *SD* | *M* | *SD* |
| No mental load | 99.47 | 1.34 | 98.89 | 2.46 | 75.56 | 14.44 |
| Low mental load | 88.82 | 19.48 | 89.65 | 10.07 | 50.33 | 25.23 |
| High mental load | 75.53 | 23.83 | 81.72 | 9.43 | 31.36 | 22.02 |
| **Miss Rate (%)** | 1-Tone paradigm |  | 2-Tone paradigm |  | 3-Tone paradigm |  |
|  | *M* | *SD* | *M* | *SD* | *M* | *SD* |
| No mental load | 0.53 | 1.34 | 1.11 | 2.46 | 24.44 | 14.44 |
| Low mental load | 11.18 | 19.48 | 10.35 | 10.07 | 49.67 | 25.23 |
| High mental load | 24.47 | 23.83 | 18.28 | 9.43 | 68.64 | 22.02 |
| **False Alarm Rate (%)** | 1-Tone paradigm |  | 2-Tone paradigm |  | 3-Tone paradigm |  |
|  | *M* | *SD* | *M* | *SD* | *M* | *SD* |
| No mental load | 0.57 | 0.37 | 0.87 | 1.02 | 9.91 | 5.00 |
| Low mental load | 4.19 | 2.90 | 5.11 | 4.70 | 9.45 | 2.84 |
| High mental load | 3.80 | 2.40 | 5.15 | 4.84 | 7.92 | 3.37 |
| **D Prime** | 1-Tone paradigm |  | 2-Tone paradigm |  | 3-Tone paradigm |  |
|  | *M* | *SD* | *M* | *SD* | *M* | *SD* |
| No mental load | 5.20 | 0.48 | 5.00 | 0.60 | 2.11 | 0.53 |
| Low mental load | 3.40 | 1.10 | 3.34 | 1.04 | 1.42 | 0.86 |
| High mental load | 2.79 | 1.13 | 2.78 | 0.76 | 0.96 | 0.83 |

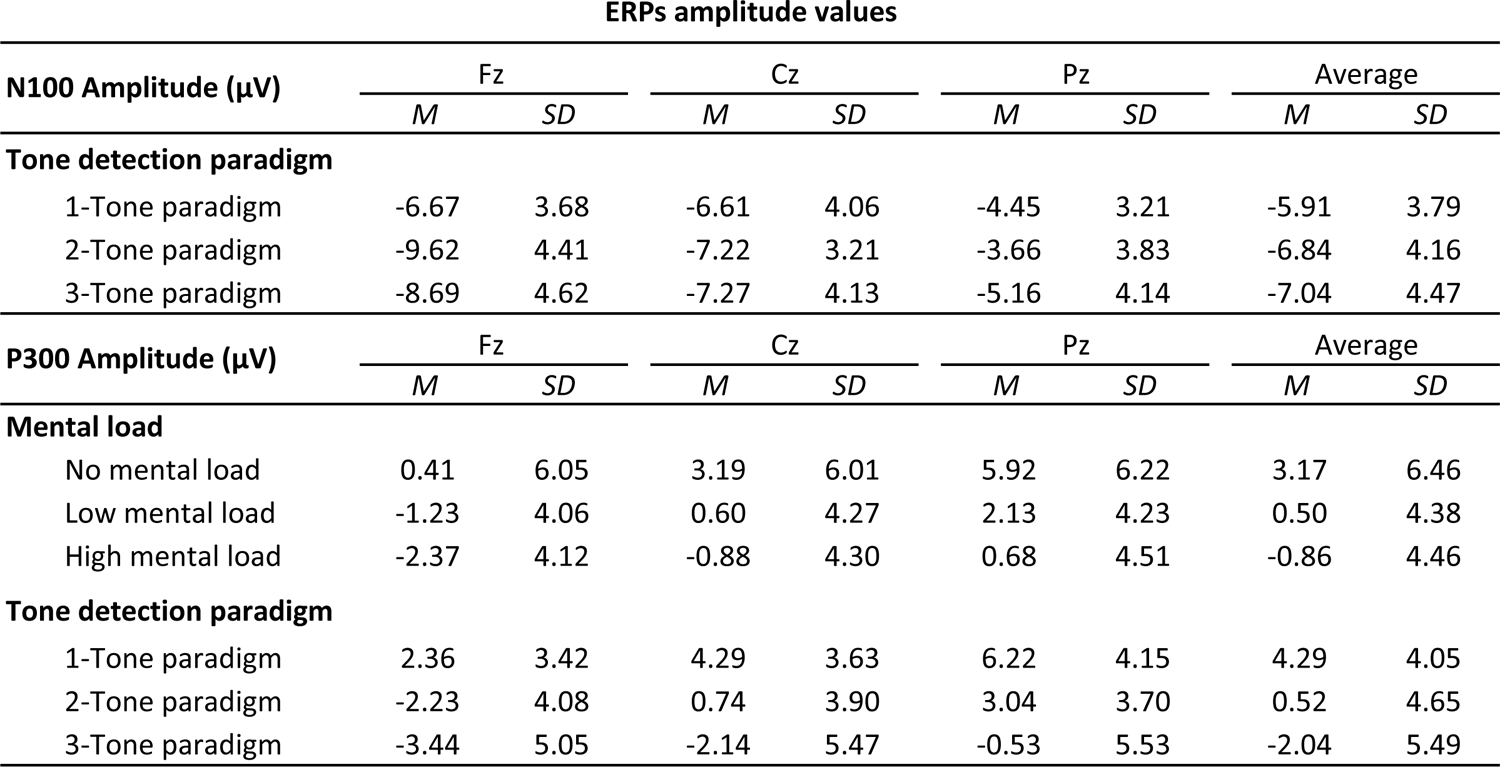

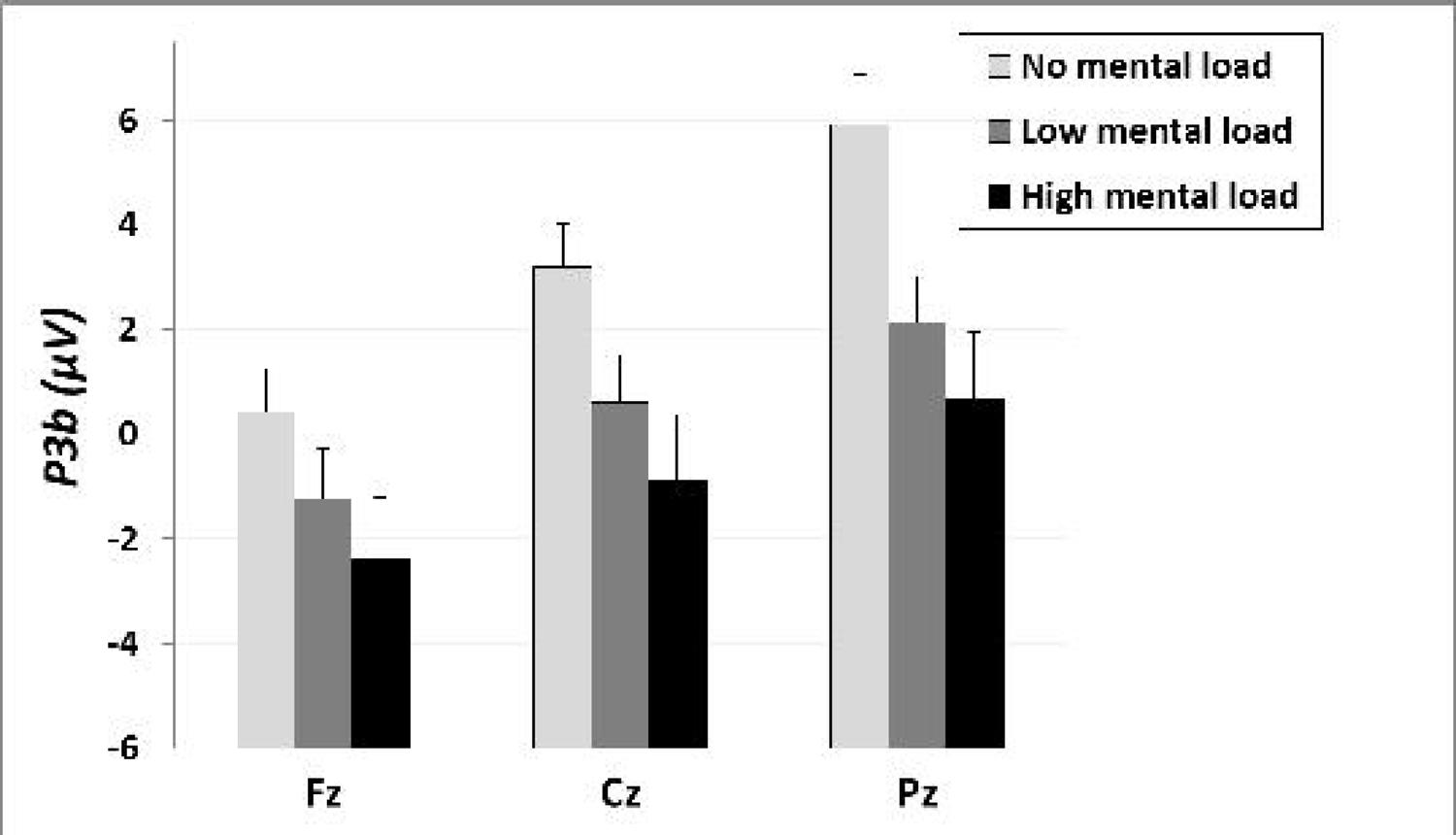

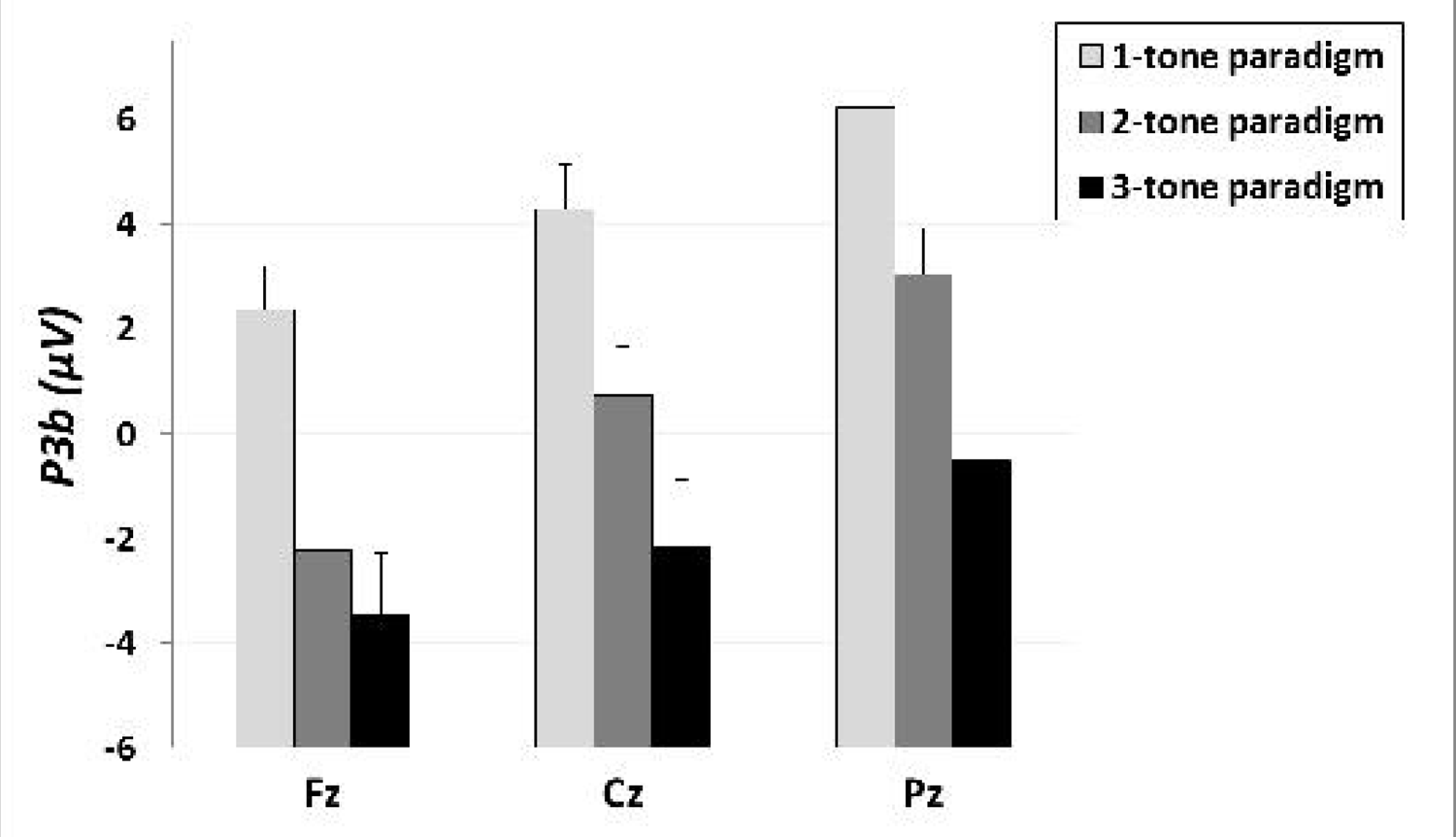

